# Potassium nutrition recover impacts on stomatal, mesophyll and biochemical limitations to photosynthesis in *Carya cathayensis* and *Hickory illinoensis*

**DOI:** 10.1101/425629

**Authors:** Chao Shen, Ruimin Huang, Yiquan Tang, Zhengjia Wang

## Abstract

Potassium (K) influences the photosynthesis process in a number of ways; However, the mechanism of photosynthetic response to the long-term supply of potassium is not yet clear. Concurrent measurements of gas exchange and chlorophyll fluorescence were made to investigate the effect of potassium nutrition on photosynthetic efficiency and stomatal conductance (gs), mesophyll conductance (gm) in Pecan (*Carya illinoensis* K.Kock) and Hickory (*Carya cathayensis* Sarg.) seedlings in a greenhouse. The results show that the photosynthetic capacity of Pecan and Hickory plants was not limited when the leaves had potassium concentrations >1.4% and 1.42% of dry weight. Most of limitation under potassium deficiency were dominated by MCL for Pecan and Hickory. Both cultivars showed remarkable improvement in S_L_, MC_L_, *J* and *V_c,max_* with additional K supplies. However, effect from potassium deficiency on photosynthesis in plant leaves was irreversible. All of S_L_, MC_L_, and B_L_ nearly half down with recovery K supply in both species. These results emphasize the important role of potassium on regulation of photosynthesis by three limitations.

## Introduction

Pecan (Hickory illinoensis K.Koch), one of the world’s efficient economic trees, can provides high quality dried fruit, good wood and other products. In China, pecan has been introduced for more than 100 years (Shi et al. 2013), and pecan have been grown in all parts of Zhejiang Province. Fruit trees are potassium sensitive crops, and their normal growth and development need adequate potassium supply. Potassium deficiency largely restricts the improvement of fruit quality. (Zhang 2016, Shen et al. 2017). However, the total potassium content of this region is low, resulting in less efficient potassium that can be absorbed and utilized by plants (Cong et al. 2016). Potassium content in plant leaves is closely related to photosynthesis (Wood et al. 2016), most studies have focused on crops (Pettigrew et al. 2010, Lu et al. 2016, Sousa et al. 2010, Li et al. 2014), while few studies have been done on fruit trees.

Potassium is one of the three microelement of plant nutrition. It is different from nitrogen and phosphorus, that exists mainly in the form of soluble inorganic salt in the cell fluid (Blevins et al. 1985), or adsorbed on the surface of the plasma colloid in the form of ions, and has no structural purpose (Tester et al. 2001). K^+^ is an activator of more than 60 enzymes in plants (Berg et al., 2010; Hu et al., 2015; Wang and Wu, 2013), including enzymes that alter carbohydrate metabolism and nitrogen metabolism, and promote protein and nucleic acid synthesis (Amtmann et al., 2008; Maathuis 1997). Potassium (such as potassium nitrate, potassium chloride and potassium citrate) is the main regulator of vacuole osmotic regulation in vacuoles (Hsiao and Läuchli 1986), regulating cell water potential and turgor pressure, thus affecting plant stomatal opening and closing movement(Jordan et al., 2008; Peiter 2011). Therefore, potassium deficiency affects various plant metabolism and osmotic adjustment, resulting in decreased leaf area, plant thin, yellow leaf wilting, ultimately inhibit the plant growth and yield formation. (Severtson et al., 2016).

K^+^ as the main regulator of guard cell permeability, its richness affects stomatal function (Shavala, 2003; Lebaudy et al. 2008; Andrés et al. 2008), thereby affecting the exchange process of the blade and outside the water and gas. Therefore, stomatal limitation was thought to be the primary cause of the decrease in leaf photosynthetic rate due to potassium deficiency (Bednarz et, al. 1998).But the research in apricot showed that leaf photosynthetic rate decreased significantly under potassium starvation (Basile et al. 2003; Oosterhuis et al. 2014; Quentin et al. 2013; Flexas et al. 2015), but the stomatal conductance is not affected, and the biochemical disorders caused by inadequate supply of potassium is the main reason that limit the photosynthetic rate (Basile et al. 2003; Flexas et al. 2012). Potassium deficiency decreased leaf chlorophyll content, decreased Rubisco activity, accelerate the generation of reactive oxygen species (ROS), may also lead to the accumulation of photosynthetic products and feedback inhibition of leaf net photosynthetic rate (Paul et al. 2003; Cakmak 2005; Araya et al., 2006; Battie-Laclau et al. 2013; Vislap et al. 2012).In recent decades, with the in-depth research on the mesophyll conductance (Bernacchi et al., 2010; Flexas et al., 2007; Galmés et al., 2007; Xiong et al., 2016), people gradually realize that potassium plays an important role in the regulation of mesophyll conductance under potassium deficiency leads to reduced mesophyll conductance on leaf photosynthetic rate limit as due to stomatal function limited (Song et al., 2011; Battie-Laclau et al., 2014).Thus, the dominant limiting factor of potassium deficiency on photosynthesis has a significant effect on the surface, this effect is based on the three factors in proportion to the size of the plant, but is regulated by internal features, including changes in the degree of stress and porosity, the resulting mesophyll layer and the physiological and biochemical characteristics.

Recently, a quantitative limitation analysis for the RuBP-limited phase of photosynthesis was proposed (Chen et al., 2013; Christian et al., 2014; Wang et al., 2015; Song et al., 2013; Wang et al., 2012). In this way, the quantitative analysis of photosynthesis limitation was conducted by combining stomatal conductance and mesophyll conductance (Song et al., 2011; Tosens et al., 2012), the total photosynthesis limitation of leaves can be divided into three components: A_L_=S_L_+M_L_+B_L_, S_L_, M_L_, B_L_, stomata, mesophyll, and biochemical limitation, respectively. To date, most studies on quantitative analysis of photosynthetic restriction have focused on crops (Sagardoy et al., 2010; Pettigrew et al., 2010; Lu et al., 2016; Sousa et al., 2010), and most of them compare genetically modified species and genotypes with large variation in leaf structure. Nevertheless, the information related to the influence of nutrient deficiency on quantitative limitation analysis for photosynthesis is missing. Previous study showed a relationship between potassium and photosynthesis in plant leaves of Pecan and Brassica napus, but do not involve the effect of potassium deficiency time on photosynthesis of leaves. For this reasons, it is desirable to develop a better understanding of the mechanisms which K supply affects photosynthesis of leaves.

## Materials and methods

### Plants materials and growth conditions

Two-year-old pecan seedlings (*Hickory illinoensis K.Koch*) were transplanted into 30.5 cm tall plastic pots with a top diameter of 25 cm, containing full-strength nutrient solution. The composition of the standard nutrient solution was as follows: 2.5 mM Ca(NO_3_)_2_, 0.5 mM Ca(H_2_PO_4_)_2_, 1.0 mMK_2_SO_4_, 0.5 mM MgSO_4_, 12.5 µM H_3_BO_3_, 1.0 µM MnSO_4_,1.0 µM ZnSO_4,_ 0.25 µM CuSO_4,_ 0.1 µM (NH_4_)_6_Mo_7_O_24_ and10 µM EDTA-Fe. The seedlings were grown in a greenhouse with natural sunlight during the day. The mean daytime maximum and minimum temperatures in the greenhouse were 28 and 20°C, with a constant relative humidity of 60%. After 2 months, the composition of the nutrient solution was altered to one of three K concentrations:0, 2.0 and 5.0 mM K, respectively. In all cases, Ca(OH)_2_ and HCl were used to adjust the pH of the nutrient solution to 5.7. The nutrient solution was changed every 7 days. All the treatments had 10 replicates with a completely random design.

### Leaf gas exchange and fluorescence measurements

Measurements were made on the youngest fully expanded leaf from 6–8 randomly selected seedlings on the 60th day of the treatment, using leaves developed after the initiation of the K nutrition treatment. Leaf gas exchange and chlorophyll fluorescence were measured simultaneously using a portable infrared gas analyser system (Li-6400, Li-Cor, Lincoln, NE, USA) equipped with an integrated leaf chamber fluorometer (Li-6400-40) at a concentration of 380 µmol mol ^−1^ CO_2_, 21% O_2_ and 50% relative humidity. Leaf chamber temperature was maintained at 28 °C. All measurements were carried out at 1200 µmol m ^−2^ s^−1^, with 90% red light and 10% blue light, which we previously determined to be just above light saturation for pecan seedlings. Once a steady state was reached (~20 min at a photosynthetic photon flux density (PPFD) of 1200 µmol m ^−2^ s^−1^), a CO_2_ response curve (*A*–*C*_i_ curve) was performed. The ambient CO_2_ concentration (*C_a_*) was lowered stepwise from 380 to 50 µmol mol−1, and then returned to 380 µmol mol−1 to re-establish the initial steady-state value of photosynthesis. *C_a_* was then increased stepwise from 380 to1800 µmol mol−1. At each *C_a_*, photosynthesis was allowed to stabilize for 3–4 min until gas exchange was steady, so that each curve was completed in 35–50 min. Corrections for the leakage of CO_2_ in and out of the Li-6400 leaf chamber, as described by Perez-Martin et al. (2009), were applied to all gas-exchange data.

The actual photochemical efficiency of photosystem II (Φ_PSII_) was determined by measuring steady-state fluorescence (*F_s_*) and maximum fluorescence during a light-saturating pulse (*F_m_*′) following the procedure of Genty et al. (1989):

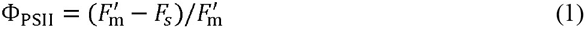

The rate of electron transport estimated from chlorophyll fluorescence is given by the equation (Bilger and Björkman1994)

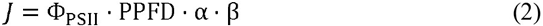

where PPFD is the photosynthetic photon flux density, α is leaf absorptance and β is the proportion of quanta absorbed by photosystem II. α⋅β was determined for each treatment from the slope of the relationship between Φ_PSII_ and Φ_CO2_ (i.e., the quantum efficiency of gross CO_2_ fixation), which was obtained by varying light intensity under non-photo respiratory conditions in an atmosphere containing <1% O_2_ (Valentini et al. 1995).

### Measurement of mitochondrial respiration rate in the light (Rd) and intercellular CO_2_ compensation point (C_i_^*^)

*R_d_* and *C_i_^*^* were determined according to the method of Laisk (1977). *A*–*C*i curves were measured using an open gas-exchange system (Li-6400, Li-Cor Inc.) equipped with an integrated light source (Li-6400-02) at three different photosynthetically active PPFDs (50, 200 and 500 mmol m^−2^ s^−1^) at six different CO_2_ levels ranging from 300 to 50 mmol CO_2_ mol^−1^ air. The curves intersected at the point where *A* is the same at different PPFDs; therefore, *A* at that point represents *R_d_*, and *C_i_* represents *C_i_^*^*.

### Estimation of gm

From combined gas-exchange and chlorophyll fluorescence measurements, the mesophyll conductance for CO_2_ (*g_m_*) was estimated according to Harley et al. (1992) as

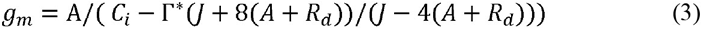

where *A*, *C*_i_, *R_d_* and *J* were determined as previously described for each treatment. Γ*^*^* is the chloroplastic CO_2_ photocompensation point calculated from the *C_i_^*^* and *R_d_* measurements according to the method of Warren et al. (2007) using a simultaneous equation with *g_m_*:

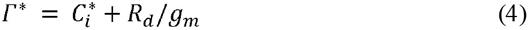

Equation (4) was then substituted into (3) and the value of *g*m was found; then Γ^*^ was calculated. The value of *Γ^*^* was found to be slightly higher for the K0-treated plants (53.9 ± 9.6 µmol mol−1), compared with the four other treatments (47.3 ± 7.5, 44.9 ± 7.8, 44.7 ± 5.1 and 44.6 ± 8.6 µmol mol^−1^ for K1, K2, K3 and K4 treatments, respectively). Changes in *Γ \** derived using the method of Laisk (1977) have been frequently observed under stress conditions such as drought (Galmés et al. 2007); therefore, we re-calculated *g*m using the non-stressed *Γ \** values (44.6 µmol mol ^−1^), which is a reasonable assumption as Γ*\** is an intrinsic property of Rubisco and thus varies only by a small amount within a species under different growing conditions. The CO_2_ concentration in the chloroplast stroma (*C_c_*) was calculated using the equation

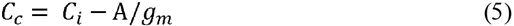

### Quantitative limitation analyses

The limitations (stomatal limitation, *S_L_*; the mesophyll conductance limitation, *MC_L_*; and the biochemical limitation, *B_L_*) imposed by K deficiency on A were investigated following Grassi and Magnani (2010). Because the fluorescence derived linear electron transport rate (J) is tightly coupled with the maximum rate of Rubisco-catalysed carboxylation (*V_c,max_*) (Galmés et al. 2007; Gallé et al. 2009), a minor modification was adopted when calculating *B_L_* using *J* instead of *V_c,max_* (Gallé et al. 2009). Relative changes in light-saturated assimilation are expressed in terms of relative changes in stomatal, mesophyll conductance and biochemical capacity as showed in Eqn 6:

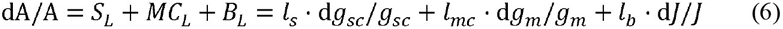

where *l_s_*, *l_mc_* and *l_b_* are the corresponding relative limitations calculated as Eqns 7–9 and *g_sc_* is stomatal conductance to CO_2_ (gs/1.6).

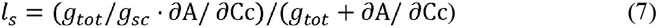

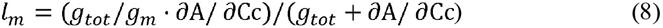

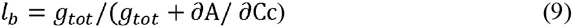

Where the gtot is the total conductance to CO_2_ from the leaf surface to carboxylation sites determined in Eqn 10. ∂*A/*∂*C_c_* was calculated as the slope of *A/C_c_* response curves over a *C_c_* range of 50–100 μmol mol1 (Tomás et al. 2012).

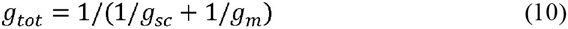

Then, the relative change of *A*, *g_sc_*, *g_m_* and *J* in Eqn 6 can be approximated by the following (Chen et al. 2013):

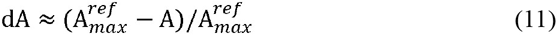

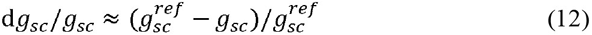

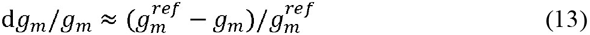

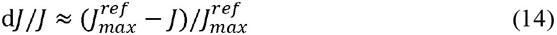

Where 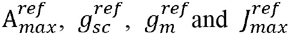are the reference values. Reference maximum values of net CO_2_ assimilation rate, stomatal and mesophyll conductance and the rate of electron transport were obtained in +K treatments; therefore, its parameters were defined as standard.

### Statistical analysis

Descriptive statistical analyses were used for the obtained parameters to assess the range of variability and standard error (SE). All data were subjected to a two-way analysis of variance (ANOVA) with SPSS 18.0 software (SPSS, Chicago, IL, USA). The difference between mean values was compared using Duncan’s multiple range test at P < 0.05. Graphics and regression analysis were performed using the GraphPad Prism 7.0 software (GraphPad, San Diego, CA).

## Results

The leaf potassium (K) concentration (%), net CO_2_ assimilation rate (*A_N_*) and chloroplastic CO_2_ concentrations (*C_C_*) of daily potassium supplied plants (control) remained mostly unchanged throughout the experiment (Fig. 1a,b,d,e), but, slightly different between species, Pecan had a little larger K and *C_C_* but lower AN compared with Hickory. After withholding potassium from plants, K and *A_N_* decreased progressively in two treatment (K0 and K2), reaching minimum values of 0.5% and < 5 μmol CO_2_ m^-2^ s^-1^, respectively, while C_C_ increased in both two cultivars, with similar trends during severe potassium stress (Fig. 3). Compare with K0 treatment, K, *A_N_* and *C_C_* of K2 recovered more quickly and closer to K5 (control, daily potassium supplied) throughout the recovery period of potassium supply. K, *A_N_* of K2 rose 41.24% and 26.98% after restoring potassium supply 7 days after in Pecan, which were significantly larger than those (18.92% and 16.16%) in Hickory.

**Fig.1.**
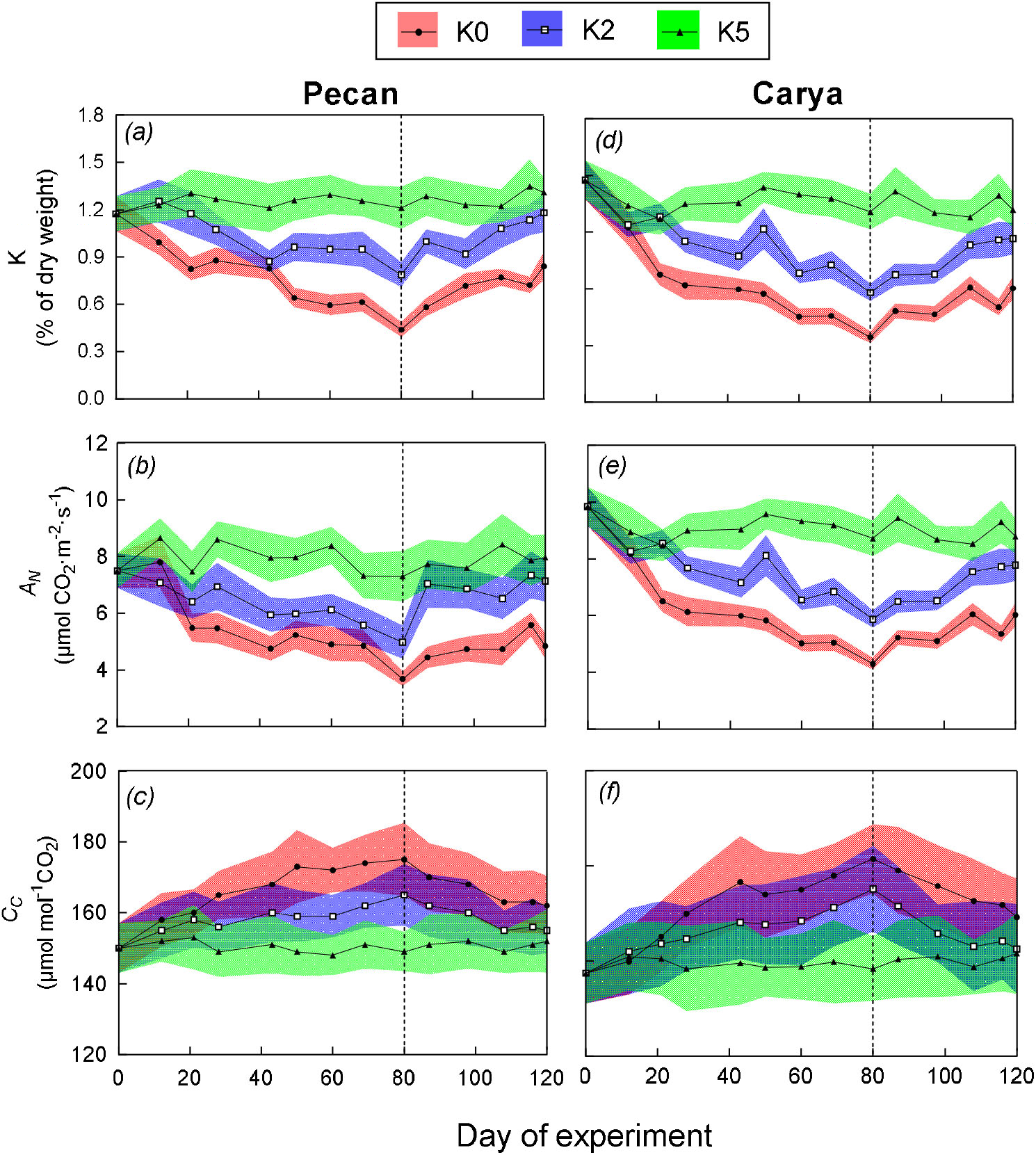
Leaf potassium percentage (%) of dry weight (K), net CO_2_ assimilation rate (*A_N_*), chloroplastic CO_2_ concentrations (*C_C_*) of Pecan and Hickory as affected by different K levels during seedling stage. The vertical dashed lines indicate the beginning of the beginning of recovering potassium supply. Data points represent means and standard errors of at least four replicates. The width of green, purple and red ribbons is the standard deviation.

After withholding potassium from plants, Mesophy ☐ conductance for CO_2_ (*g_m_*), stomatal conductance for CO_2_ (*g_s_*), electron transport rate (*J*) and maximum velocity of carboxylation (*V_c,max_*) decreased progressively in all treatments of Pecan and Hickory, reaching minimum values of 0.0265 mol CO_2_m^-2^s^-1^, 0.0457 molm^-2^s^-1^, 99.94μmol e^-1^m^-2^s^-1^ and 47.5904μmol CO_2_m^-2^s^-1^ after severe potassium stress for 80d, respectively. After recovering potassium supply, the large restoration of *g_m_*, *g_s_*, *J* and *V_c,ma_*_x_ were observed in the K0 and K2 treatments in two cultivations. The gm of K0 and K2 recovered 55.16% and 71.54% to control values (0.09684 mol CO_2_m^-2^s^-1^) of Pecan (Fig. 2a), respectively; gm of K0 and K2 recovered 55.65% and 81.93% to control values(0.09938 mol CO_2_m^-2^s^-1^) of Hickory (Fig. 2b), respectively; gs recovered 76.80% and 90.64% to control values (0.1082 molm^-2^s^-1^) of Hickory (Fig. 2f) respectively, which were nearly 10 percentage points more than of Pecan. The restoration of *J* and *V_c,max_* under both K0 and K2 treatments were reached to control values after recovering potassium supply for 40d (Fig. 2c, d, g h), respectively. However, slightly different levels of recovery time of *g_m_ g_s_* and *V_c,max_* were observed in K2 treatments, with the three photosynthetic parameters show faster recovery speed after recovering potassium supply. Under K5 treatment, *g_m_*, *g_s_*, *J* and *V_c,max_* of Pecan and Hickory almost unaltered throughout the experiment, while those photosynthetic parameters of Hickory were slightly larger than Pecan.

**Fig.2.**
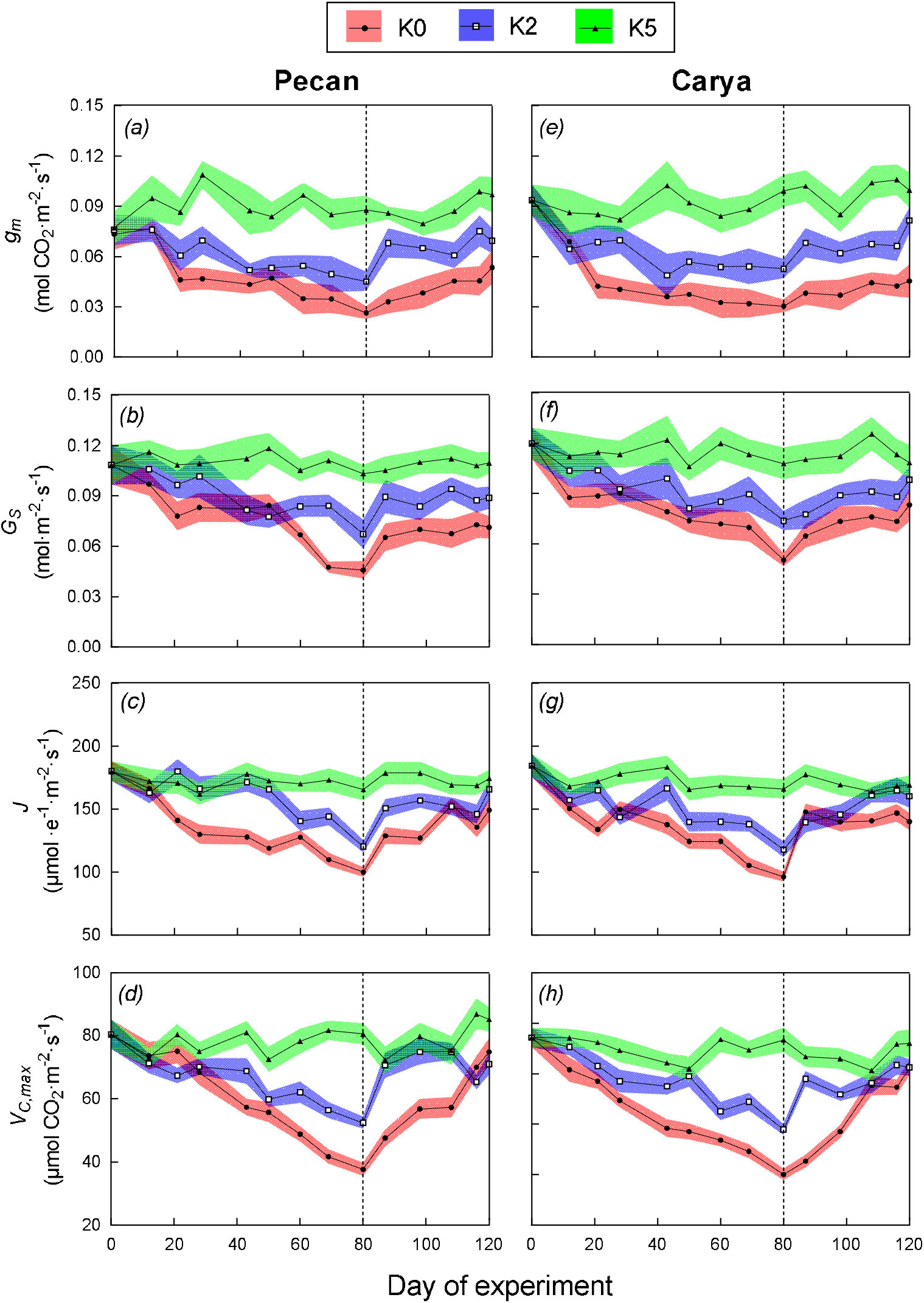
Mesophy ☐ conductance for CO_2_ (*g_m_*), stomatal conductance for CO_2_ (*g_s_*), electron transport rate (*J*) and maximum velocity of carboxylation (*V_c,max_*) of Pecan and Hickory as affected by different K levels during seedling stage. The vertical dashed lines indicate the beginning of the beginning of recovering potassium supply. Data points represent means and standard errors of at least four replicates. The width of green, purple and red ribbons is the standard deviation.

When plotting all K content (consisting all of K5, K2 and K0 under potassium stress and recover, respectively) against the corresponding calculated *A_N_*, *J*, *g_s_* and *g_m_*, highly significant positive correlation relationships were obtained pooling potassium supply and potassium stress data together, although two different functions were derived for the Pecan and Hickory (Fig. 3a-d). In these four figures, slightly steep slope was determined for the Pecan data set, however, less clear difference was observed between two cultivars. Moreover, K content on *A_N_*, *J*, *g_s_* and *g_m_* values resulted in almost similar slopes of linear regression for both two cultivars, respectively.

**Fig.3.**
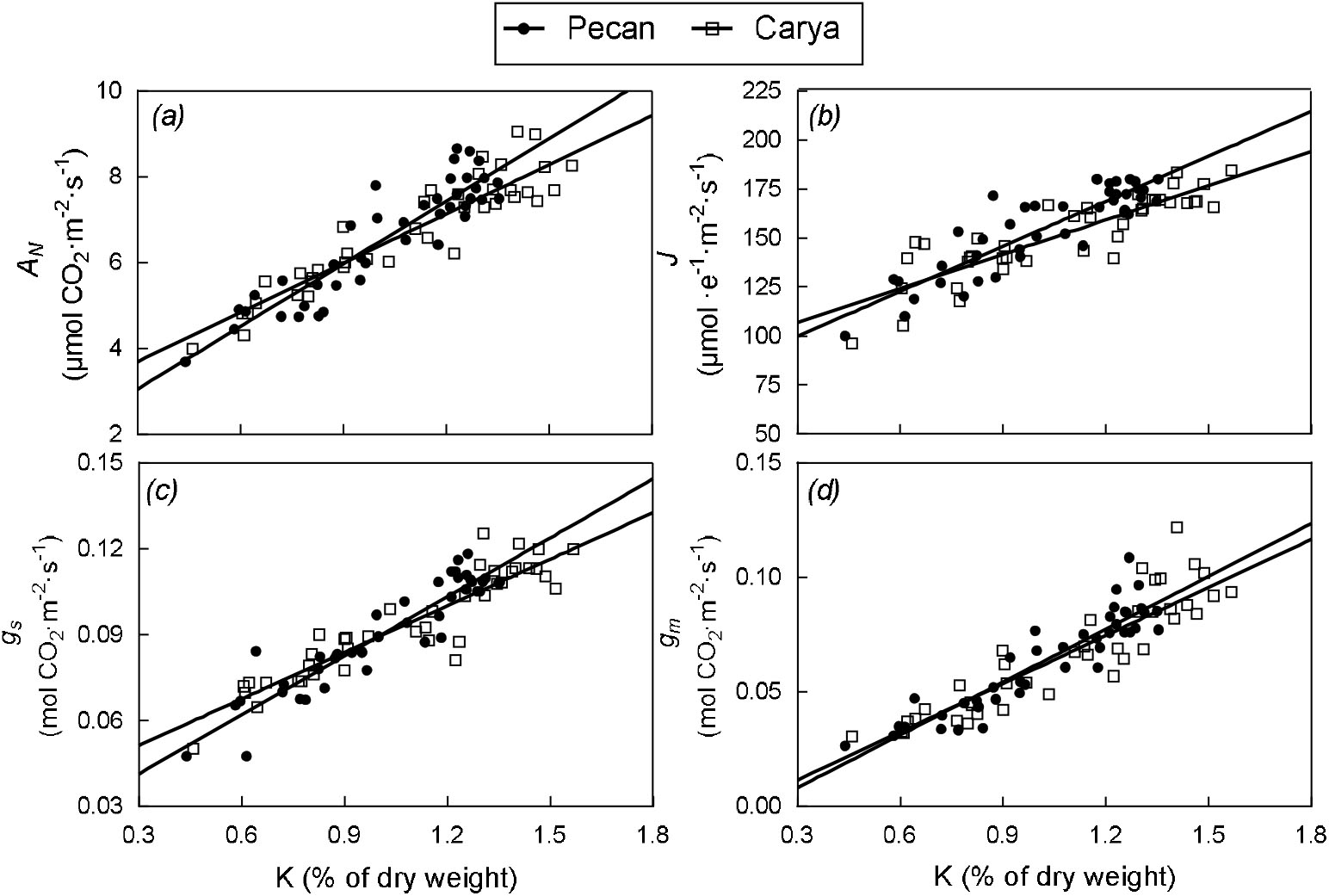
The relationships between leaf potassium percentage(%) of dry weight (K) and net CO_2_ assimilation rate (*A_N_*), electron transport rates (*J*), stomatal conductance for CO_2_ (*g_s_*) and mesophy ☐ conductance for CO_2_ (*g_m_*) derived from data of the whole experimental periods. Circles and diamonds denote Pecan and Hickory data, respectively. Data points represent means and standard errors of at least four replicates.

The A_N_ correlated negatively with chloroplastic CO_2_ concentrations (*C_C_*), while, A_N_ in Hickory was slightly higher Pecan. When plotting all *A_N_* (consisting all of K5, K2 and K0 under potassium stress and recover potassium, respectively against the corresponding calculated *J* and gm, highly significant positive correlation relationships were obtained pooling potassium supply and potassium stress data together, although two different functions were derived for the Pecan and Hickory (Fig. 4b,c).In addition, *A_N_* on *J* and gm values resulted in almost similar slopes of linear regression for both two cultivars, respectively.

**Fig.4.**
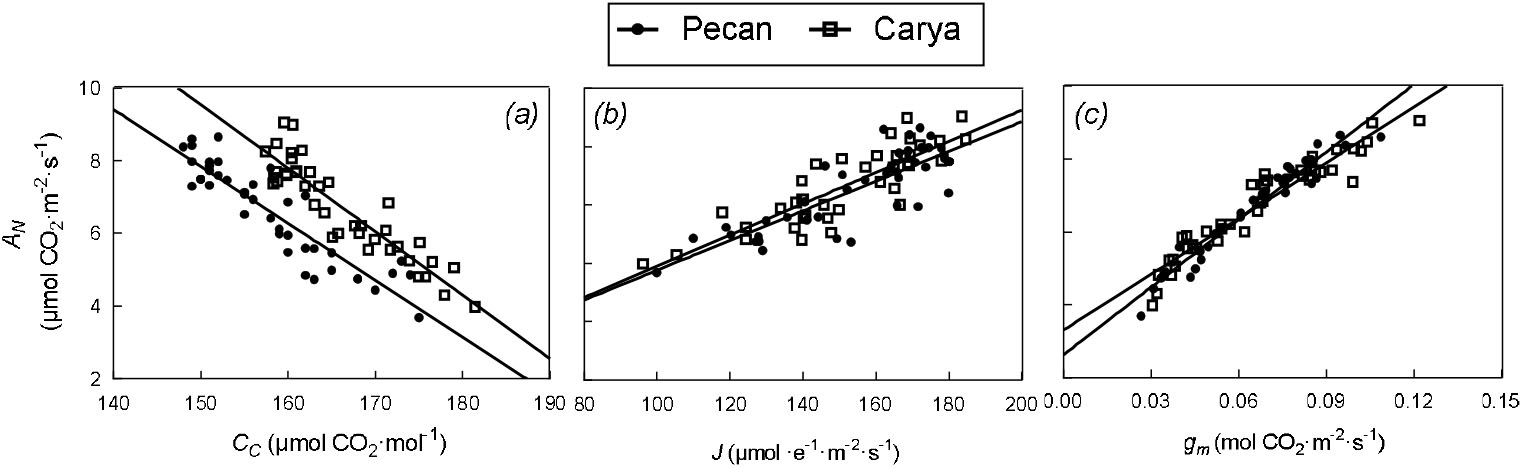
The relationships between leaf potassium percentage (%) of dry weight (K) and net CO_2_ assimilation rate (*A_N_*), electron transport rates (*J*), stomatal conductance for CO_2_ (*g_s_*) and mesophy ☐ conductance for CO_2_ (*g_m_*) derived from data of the whole experimental periods. Circles and diamonds denote Pecan and Hickory data, respectively. Data points represent means and standard errors of at least four replicates.

When analyzing the effects of potassium on photosynthesis different indicators of stress intensity can be used. In order to enhance the comparability of our data with other experiment, we, therefore expressed relative limitations in terms of both of both K (% of dry weight) (Fig. 5)and day of experiment (Fig. 6).what the stress index adopted, small differences between Pecan and Hickory. With increasing potassium stress intensity MC_L_, B_L_ and S_L_ increased significantly, and, the increase rate of MC_L_ is greater than that of B_L_ and S_L_. At mild-to-moderate potassium stress levels (corresponding to values of K > 0.9% of dry weight), about half of the decline in A_N_ was attributable to mesophy ☐ resistance.

**Fig.5.**
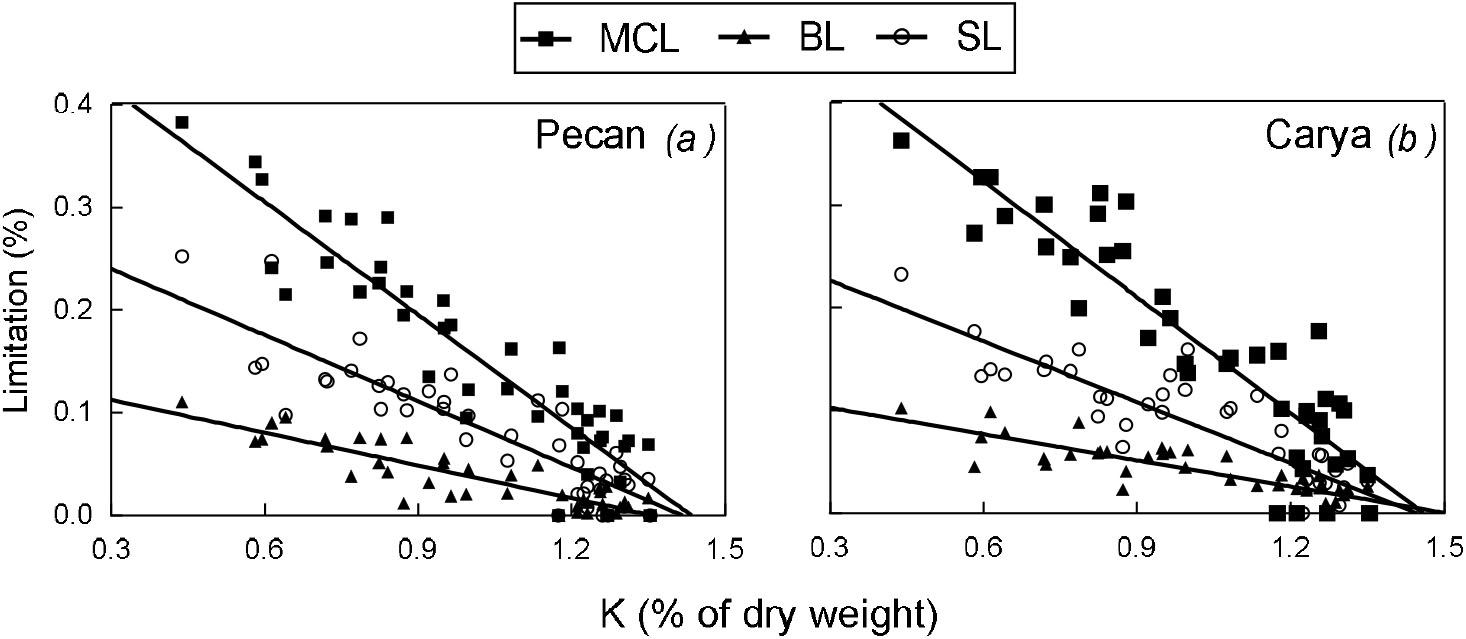
The relationships between photosynthetic limitations and leaf potassium percentage (%) of dry weight (K) of Pecan (a) and Hickory (b).S_L_ MC_L_ B_L_ denote stomatal, mesophy ☐ conductance and biochemical limitation respectively. Each point in the same shape represents a calculation (42 values were calculated for each limitation)

**Fig.6.**
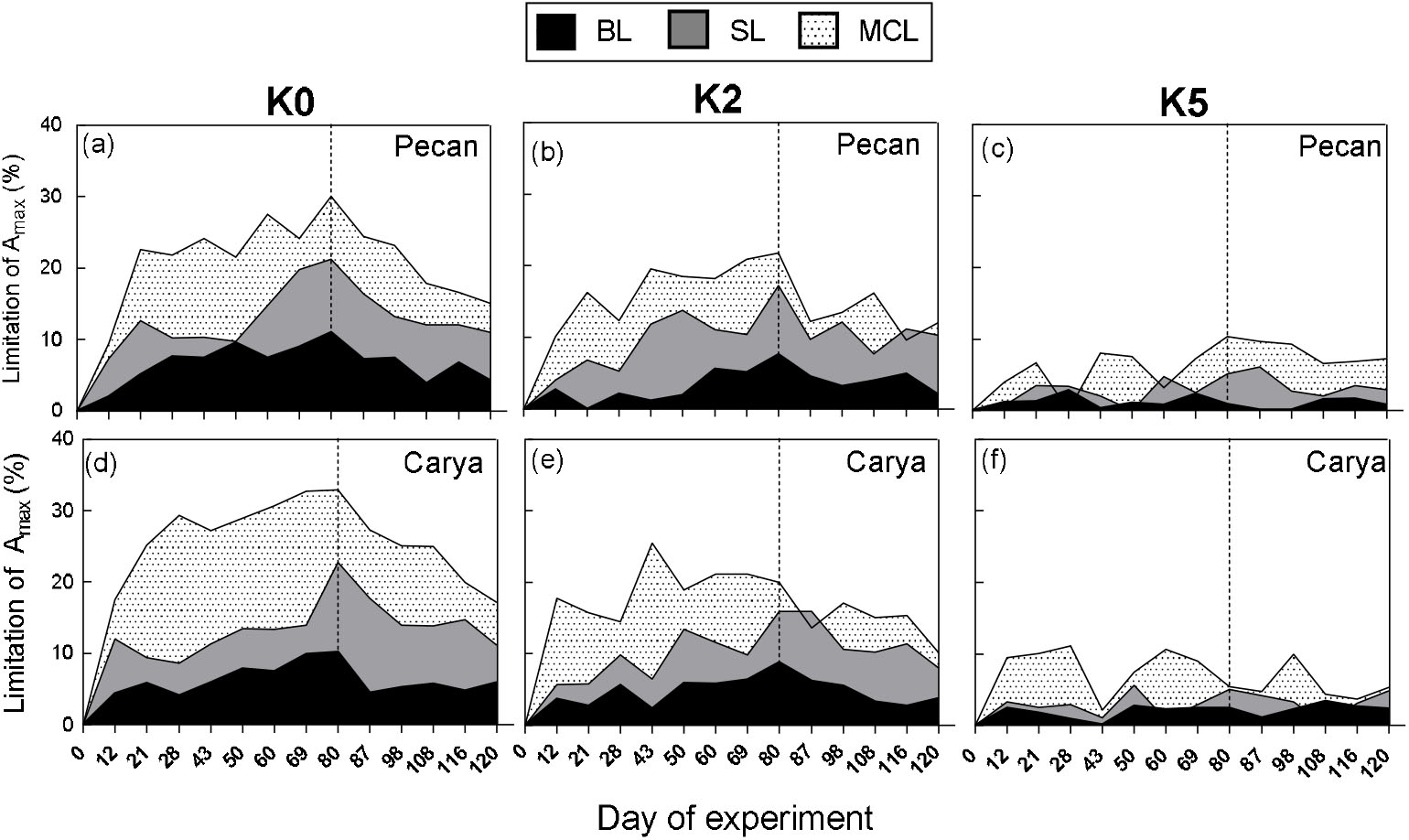
Quantitative limitation analysis of photosynthetic CO_2_ assimilation during potassium stress and subsequently recovering. The shaded areas represent the percentage of stomatal (SL), mesophyll (M_L_), and biochemical (B_L_) limitation based on control values of *A_N_*, *g_s_*, *g_m_*, and *V_c,max_* and *J*. Calculations were done with mean values of at least five measurements per treatment and day. The vertical dashed lines indicate the beginning of the beginning of recovering potassium supply. Data points represent means and standard errors of at least four replicates.

Quantitative limitation analysis of photosynthesis underlined the above-described changes during potassium stress and recovery after subsequent recover potassium. In all three experiments, MC_L_ play a major role of the total limitation under severe potassium stress. In K0 and K2 treatments, MC_L_ made up > 40% and >50% of the total limitation under severe potassium stress in Pecan and Hickory, while S_L_ accounted for only up to 20%. Furthermore, B_L_ did not exceed 10% of the total limitation. As already observed for the *A_N_*, *g_s_*, and gm data during stress and recovery, almost no limitation of S_L_ and B_L_ during potassium stress and after recover potassium (Fig. 6c,f). Limitation of photosynthetic recovery of the K0 and K2 plants was mainly driven by a still high MC_L_ and somewhat lower S_L_ and B_L_ (Fig. 5a, b, d, e).The delayed recovery of photosynthesis in the K0 plants was mainly due to a maintained high proportion of MCL and SL (Fig. 5d) during several days of recover potassium (Fig. 6a, d), while SL contributed only partially to the total limitation in the initial phase of re-watering. The recovery of photosynthesis in the K2 plants is due to a rapidly decreasing MCL during the recover potassium period, while SL is nearly equaled MCL in the later phase of recover potassium (Fig. 6b, e).

## Discussion

In the present study, the same experimental design was applied to Brassica napus under four different potassium treatments, followed by a potassium stress period and a recover after recover potassium. The most important difference between the four potassium treatments was observed: the rate of photosynthetic recovery was the slowest under K0 treatment and the quickest under K2 treatment of Pecan. Then, extremely low photosynthetic limitation in plant leaves of Pecan and Hickory under K5 treatment. A positive relationship between K supply and *A_N_* had reported in numerous previous studies. However, photosynthesis rates may not have changed as a response to K treatment due to relatively higher leaf K concentration, which were far more than the values (1.04% and 1.28 % of dry weight) proposed by Gierth (2007) and Zhifeng Lu (2016) respectively, as are the values (1.4 % and 1.42%) in this study.

Withholding potassium resulted in a closure of stomata, which was accompanied by a marked decrease of net photosynthesis (*A_N_*) in K0 and K2 treatment of Pecan and Hickory (Fig. 2). Throughout periods of potassium stress imposition, stomatal conductance (*g_s_*) and AN followed the same course, indicating a strong correlation between them, which has been shown elsewhere (Medrano et al., 2002). Moreover, plants under K5 treatment displayed a inequable course for stomatal conductance (*g_s_*) of Pecan and Hickory. In addition, mesophyll conductance (*g_m_*) showed a similar trend with stomatal conductance (*g_s_*) under all three potassium supply in Pecan and Hickory. These results are in line with previous studies, where a decrease of gm has been observed during potassium stress (Song et al., 2011; Lu et al., 2016; Wang et al., 2012; Battie-Laclau et al., 2014). The rapid restoration of *J* and *V_cmax_* to control values during prolonged recover potassium, while the restoration of *g_s_* and *g_m_* is could not reach the level of the control value, especially under K0 stress of Pecan and Hickory. These results are indicating restored *J* and *V_cmax_* during prolonged recover potassium presumably facilitated photosynthetic recovery after the beginning of re-watering, because *A_N_* and *J*, *V_cmax_* were immediately restored to control values within the initial phase.

Lots of previous studies have shown total photosynthetic dramatically decreased with decreasing K supply. (Bednarz et al.1998; Jin et al. 2011; Wang et al. 2012; Battie-laclau et al. 2014; erel et al. 2015). And the results of quantitative analysis revealed that the three components contributing to total photosynthetic limitation, namely, S_L_, MC_L_, and B_L_ varied at varying K concentrations. The decline of *P_n_* with the decrease of potassium concentration in leaves, while all of S_L_, B_L_ and MC_L_ were decreased, B_L_ was markedly lower and MC_L_ was higher, especially at lower treatment in both cultivars (Fig. 6). This can be attributed to prioritization of allocation of excess K to the cytosol for metabolic activity rather than to reduce the transmission resistance of CO_2_ in the chloroplast. (Reich et al., 1997; Pettigrew 1999; Zhao et al., 2001; Battie-Laclau et al., 2013; Tomás et al., 2016). Then our quantitative limitation analysis showed that SL was always higher than B_L_ under three treatments in both species. This pattern of response is consistent with that decried by other authors (Tanaka et al., 2005; Christian et al., 2014; Peiter et al., 2011). Although several studies have reported that stomatal closure (S_L_) plays by far the main role in the decline of photosynthesis, even at rather severe levels of potassium stress (Bednarz 1998; shavala, 2003; Lebaudy, 208; Andrés, 2014), but our study draw different conclusions that MC_L_ is the dominant factor in the AN reduction rather than the S_L_ in both species. This striking discrepancy is explained by the fact that potassium stress increases the transmission distance of CO_2_ in the chloroplast (Zhao et al., 2001; Battie-Laclau et al., 2014).

An even clear picture of the potassium effects emerges when the different limitations are plotted against potassium stress and subsequent recovering (Fig. 6). In both species, all the limitations increased with increasing potassium stress and duration of treatment, but their relative contribution changed. Moreover, in the early stage of potassium treatment (0-21d), all the three limitations under k0 and K2 rapidly risen. In the middle and late period (21-80d), all the limitations maintained their high level and changed little under K0 treatment, this could be the acclimation of plants to severe potassium deficiency, a similar situation had been appeared in the plant drought test (Galle et al., 2009). After recover potassium, obvious decline could be observed of all the three limitations under K0 and K2 treatment in both species. However, S_L_ and MC_L_ were not hopely recovered to the control level (K5), they remain at a high level until the end of the experiment. The most likely explanation for this discrepancy might be derived from the effect of potassium stress on the irreversible structural effects of plant leaves. In this sense, potassium deficiency induced the increase of with leaf dry mass per area (MA)(Reich et al., 1997; Pettigrew, 1999; Zhao et al., 2001; Song et al., 2011), reducing the leaves thickness and the volume of cell gap in the leaves, thus reducing the gas conduction ability of CO_2_ (Battie-Laclau et al., 2014).

In conclusion, A_N_ of Pecan and Craya plants declined by increasing S_L_, MC_L_, and B_L_ under prolonged severe potassium stress. Pecan, needed K (1.4% of dry weight less than that of Hickory) to avoid the decline of A_N_. All of S_L_, MC_L_, and B_L_ had a sharp decline under K0 and K5treatment in both species. In summary, the present study strongly reinforces the important role of *g_s_*, *g_m_*, *J* and *V_c,max_* during recovery from potassium stress induced inhibition of photosynthesis, and shows for the first time that such a role depends on the Long-term processing.

## References

Amtmann, A., Troufflard, S., & Armengaud, P. (2010). The effect of potassium nutrition on pest and disease resistance in plants. Physiologia Plantarum, 133(4), 682–691.

Andrés, Z., Pérez-Hormaeche, J., Leidi, E. O., Schlücking, K., Steinhorst, L., & Mclachlan, D. H., et al. (2014). Control of vacuolar dynamics and regulation of stomatal aperture by tonoplast potassium uptake. Proceedings of the National Academy of Sciences of the United States of America, 111(17), 1806–14.

Araya, T., Noguchi, K., & Terashima, I. (2006). Effects of carbohydrate accumulation on photosynthesis differ between sink and source leaves of phaseolus vulgaris l. Plant & Cell Physiology, 47(5), 644–52.

Basile, B., Reidel, E. J., Weinbaum, S. A., & Dejong, T. M. (2003). Leaf potassium concentration, co 2, exchange and light interception in almond trees (prunus dulcis, (mill) d.a. webb). Scientia Horticulturae, 98(2), 185–194.

Battie‐Laclau, P., Laclau, J., Domec, J., Christina, M., Bouillet, J., & Cassia Piccolo, M., et al. (2014). Effects of potassium and sodium supply on drought‐adaptive mechanisms in eucalyptus grandis plantations. New Phytologist, 203(2), 401–413.

Battie‐Laclau, P., Laclau, J., Beri, C., Mietton, L., Muniz, M. R. A., & Arenque, B. C., et al. (2013). Photosynthetic and anatomical responses of eucalyptus grandis leaves to potassium and sodium supply in a field experiment. Plant Cell & Environment, 37(1), 70–81.

Bednarz, C. W., Oosterhuis, D. M., & Evans, R. D. (1998). Leaf photosynthesis and carbon isotope discrimination of cotton in response to potassium deficiency. Environmental & Experimental Botany, 39(2), 131–139.

Berg, W. K., Cunningham, S. M., Brouder, S. M., Joern, B. C., Johnson, K. D., & Volenec, J. J.(2010). Influence of phosphorus and potassium on alfalfa yield, taproot c and n pools, and transcript levels of key genes after defoliation. Crop Science, 49(3), 974–982.

Blevins, D. G. (1985). Role of potassium in protein metabolism in plants.

Bernacchi, C. J., Singsaas, E. L., Pimentel, C., Jr, A. R. P., & Long, S. P. (2010). Improved temperature response functions for models of rubisco‐limited photosynthesis. Plant Cell & Environment, 24(2), 253–259.

Bilger, W., & Björkman, O. (1994). Relationships among violaxanthin deepoxidation, thylakoid membrane conformation, and nonphotochemical chlorophyll fluorescence quenching in leaves of cotton (gossypium hirsutum, l.). Planta, 193(2), 238–246.

Cakmak, I. (2005). Role of potassium in alleviating the detrimental effects of abiotic stress plants. j plant nutr soil sci. Journal of Plant Nutrition & Soil Science, 168(4), 521–530.

Chen, T. W., Henke, M., Kahlen, K., Visser, P. H. B. D., Buck-Sorlin, G., & Wiechers, D., et al.(2013). Revealing the relative importance of photosynthetic limitations in cucumber canopy. Fspm.

Christian Zörb, Mehmet Senbayram, & Edgar Peiter. (2014). Potassium in agriculture – status and perspectives ☆. Journal of Plant Physiology, 171(9), 656–669.

Cong, R., Li, H., Zhang, Z., Ren, T., Li, X., & Lu, J. (2016). Evaluate regional potassium fertilization strategy of winter oilseed rape under intensive cropping systems: large-scale field experiment analysis. Field Crops Research, 193, 34–42.

Erel, R., Yermiyahu, U., Ben-Gal, A., Dag, A., Shapira, O., & Schwartz, A. (2015). Modification of non-stomatal limitation and photoprotection due to k and na nutrition of olive trees. Journal of Plant Physiology, 177, 1–10.

Flexas, J., & Diaz-Espejo, A. (2015). Interspecific differences in temperature response of mesophyll conductance: food for thought on its origin and regulation. Plant, cell & environment, 38(4), 625–8.

Flexas, J., Barbour, M. M., Brendel, O., Cabrera, H. M., Carriquí, M., & Díaz-Espejo, A., et al.(2012). Mesophyll diffusion conductance to co 2 : an unappreciated central player in photosynthesis. Plant Science An International Journal of Experimental Plant Biology, s193–194(1), 70–84.

Flexas, J., Díazespejo, A., Berry, J. A., Cifre, J., Galmés, J., & Kaldenhoff, R., et al. (2007). Analysis of leakage in irga’s leaf chambers of open gas exchange systems: quantification and its effects in photosynthesis parameterization. Journal of Experimental Botany, 58(6), 1533.

Galle, A., & Flexas, J. (2009). The role of mesophyll conductance during water stress and recovery in tobacco (nicotiana sylvestris): acclimation or limitation?. Journal of Experimental Botany, 60(8), 2379–2390.

Galmes, J., & Medrano, H. J. (2007). Photosynthetic limitations in response to water stress and recovery in mediterranean plants with different growth forms. New Phytologist, 175(1), 81–93.

Genty, B., Briantais, J. M., & Baker, N. R. (1989). The relationship between the quantum yield of photosynthetic electron transport and quenching of chlorophyll fluorescence. BBA - General Subjects, 990(1), 87–92.

Gierth, M., & Mäser, P. (2007). Potassium transporters in plants – involvement in k+ acquisition, redistribution and homeostasis. Febs Letters, 581(12), 2348–2356.

Grassi, G., & Magnani, F. (2010). Stomatal, mesophyll conductance and biochemical limitations to photosynthesis as affected by drought and leaf ontogeny in ash and oak trees. Plant Cell & Environment, 28(7), 834–849.

Harley, P. C., Loreto, F., Di, M. G., & Sharkey, T. D. (1992). Theoretical considerations when estimating the mesophyll conductance to co(2) flux by analysis of the response of photosynthesis to co(2). Plant Physiology, 98(4), 1429–1436.

Hsiao, T. C., & Läuchli, A. (1986). Role of potassium in plant-water relations. Advances in Plant Nutrition.

Hu, W., Yang, J., Meng, Y., Wang, Y., Chen, B., & Zhao, W., et al. (2015). Potassium application affects carbohydrate metabolism in the leaf subtending the cotton (gossypium hirsutum, l.) boll and its relationship with boll biomass. Field Crops Research, 179, 120–131.

Jordan-Meille, L., & Pellerin, S. (2008). Shoot and root growth of hydroponic maize (zea mays l.) as influenced by k deficiency. Plant & Soil, 304(1-2), 157–168.

Lai sk, A. Kh. (Agu Khei novich). (1977). Kinetics of photosynthesis and photorespiration of c3 in plants. Mccarthy.

Lebaudy, A., Vavasseur, A., Hosy, E., Dreyer, I., Leonhardt, N., & Thibaud, J. B., et al. (2008). Plant adaptation to fluctuating environment and biomass production are strongly dependent on guard cell potassium channels. Proceedings of the National Academy of Sciences of the United States of America, 105(13), 5271–5276.

Li, X., Chunsheng, M. U., Lin, J., Wang, Y., & Xiujun, L. I. (2014). Effect of alkaline potassium and sodium salts on growth, photosynthesis, ions absorption and solutes synthesis of wheat seedlings. Experimental Agriculture, 50(1), 144–157.

Lu, Z., Lu, J., Pan, Y., Lu, P., Li, X., & Cong, R., et al. (2016). Anatomical variation of mesophyll conductance under potassium deficiency has a vital role in determining leaf photosynthesis. Plant Cell & Environment, 39(11), 2428–2439.

Lu Z, Ren T, Pan Y, Li X, Cong R, & Lu J. (2016). Differences on photosynthetic limitations between leaf margins and leaf centers under potassium deficiency for brassica napus l. Scientific Reports, 6, 21725.

Maathuis, F. J., Ichida, A. M., Sanders, D., & Schroeder, J. I. (1997). Roles of higher plant k^+^ channels. Plant Physiology, 114(4), 1141–1149.

Medrano, H., Escalona, J. M., Bota, J., Gulías, J., & Flexas, J. (2002). Regulation of photosynthesis of c3 plants in response to progressive drought: stomatal conductance as a reference parameter. Annals of Botany, 89(7), 895–905.

Oosterhuis, D. M., Loka, D. A., Kawakami, E. M., & Pettigrew, W. T. (2014). The physiology of potassium in crop production. Advances in Agronomy, 126, 203–233.

Paul, M. J., & Pellny, T. K. (2003). Carbon metabolite feedback regulation of leaf photosynthesis and development. Journal of Experimental Botany, 54(382), 539–547.

Peiter, E. (2011). The plant vacuole: emitter and receiver of calcium signals. Cell Calcium, 50(2), 120–128.

Perezmartin, A., Flexas, J., Ribascarbó, M., Bota, J., Tomás, M., & Infante, J. M., et al. (2009). Interactive effects of soil water deficit and air vapour pressure deficit on mesophyll conductance to co2 in vitis vinifera and olea europaea. Journal of Experimental Botany, 60(8), 2391.

Pettigrew, W. T. (1999). Potassium deficiency increases specific leaf weights and leaf glucose levels in field-grown cotton. Agronomy Journal, 91(6), 962–968.

Pettigrew, W. T. (2010). Potassium influences on yield and quality production for maize, wheat, soybean and cotton. Physiologia Plantarum, 133(4), 670–681.

Quentin, A. G., Close, D. C., Hennen, L. M. H. P., & Pinkard, E. A. (2013). Down-regulation of photosynthesis following girdling, but contrasting effects on fruit set and retention, in two sweet cherry cultivars. Plant Physiology & Biochemistry Ppb, 73(73C), 359–367.

Reich, P. B., Walters, M. B., & Ellsworth, D. S. (1997). From tropics to tundra: global convergence in plant functioning. Proceedings of the National Academy of Sciences of the United States of America, 94(25), 13730–13734.

Sagardoy, R., Vázquez, S., Florezsarasa, I. D., Albacete, A., Ribascarbó, M., & Flexas, J., et al. (2010). Stomatal and mesophyll conductances to co2 are the main limitations to photosynthesis in sugar beet (beta vulgaris) plants grown with excess zinc. New Phytologist, 187(1), 145–158.

Severtson, D., Callow, N., Flower, K., Neuhaus, A., Olejnik, M., & Nansen, C. (2016). Unmanned aerial vehicle canopy reflectance data detects potassium deficiency and green peach aphid susceptibility in canola. Precision Agriculture, 17(6), 1–19.

Shabala, S. (2003). Regulation of potassium transport in leaves: from molecular to tissue level. Annals of Botany, 92(5), 627–634.

Shen, C., Wang, J., Jin, X., Liu, N., Fan, X., & Dong, C., et al. (2017). Potassium enhances the sugar assimilation in leaves and fruit by regulating the expression of key genes involved in sugar metabolism of asian pears. Plant Growth Regulation(1), 1–14.

Shi JJ, Ye SY, Yu SQ, Wang ZJ (2013). Genetic diversity SSR analysis of 37 newly introduced hickory hickory varieties. Journal of Anhui Agricultural University. 40(1):42–6.

Song, H. J., Jian, Q. H., Xue, Q. L., Bing, S. Z., Jia, S. W., & Zheng, J. W., et al. (2011). Effects of potassium supply on limitations of photosynthesis by mesophyll diffusion conductance in carya cathayensis. Tree Physiology, 31(10), 1142.

Song, Q., Zhang, G., & Zhu, X. G. (2013). Optimal crop canopy architecture to maximise canopy photosynthetic co2 uptake under elevated co2 – a theoretical study using a mechanistic model of canopy photosynthesis. Functional Plant Biology Fpb, 40(2), 109–124.

Sousa, J. V. D., Rodrigues, C. R., Luz, J. M. Q., Carvalho, P. C. D., Rodrigues, T. M., & Brito, C. H. D. (2010). Foliar application of the potassium silicate in corn: photosynthesis, growth and yield. Bioscience Journal, 26(4), 502–513.

Tanaka, Y., Sano, T., Tamaoki, M., Nakajima, N., Kondo, N., & Hasezawa, S. (2005). Ethylene inhibits abscisic acid-induced stomatal closure in arabidopsis thaliana. Plant Physiology, 138(4), 2337–2343.

Tester, M., & Leigh, R. A. (2001). Partitioning of nutrient transport processes in roots. Journal of Experimental Botany, 52(356), 653–653.

Tosens, T., Ülo Niinemets, Westoby, M., & Wright, I. J. (2012). Anatomical basis of variation in mesophyll resistance in eastern australian sclerophylls: news of a long and winding path. Journal of Experimental Botany, 63(14), 5105–5119.

Tosens, T., Nishida, K., Gago, J., Coopman, R. E., Cabrera, H. M., & Carriquí, M., et al. (2016). The photosynthetic capacity in 35 ferns and fern allies: mesophyll co2 diffusion as a key trait. New Phytologist, 209(4), 1576–1590.

Valentini, R., Epron, D., Angelis, P. D., Matteucci, G., & Dreyer, E. (2010). In situ estimation of net co2 assimilation, photosynthetic electron flow and photorespiration in turkey oak (q. cerris l.) leaves: diurnal cycles under different levels of water supply. Plant Cell & Environment, 18(6), 631–640.

Vislap, V. (2012). Developmental changes in mesophyll diffusion conductance and photosynthetic capacity under different light and water availabilities in populus tremula: how structure constrains function. Plant Cell & Environment, 35(5), 839–856.

Wang, M., Zheng, Q., Shen, Q., & Guo, S. (2013). The critical role of potassium in plant stress response. International Journal of Molecular Sciences, 14(4), 7370–7390.

Wang, N., Hua, H., Eneji, A. E., Li, Z., Duan, L., & Tian, X. (2012). Genotypic variations in photosynthetic and physiological adjustment to potassium deficiency in cotton (gossypium hirsutum). Journal of Photochemistry & Photobiology B Biology, 110(9), 1–8.

Wang, Y., & Wu, W. H. (2015). Genetic approaches for improvement of the crop potassium acquisition and utilization efficiency. Current Opinion in Plant Biology, 25, 46–52.

Wood, R., & Parish, M. (2003). The mechanisms and viticultural factors governing potassium accumulation in the grape berry - part 1.

Xiong, D., Liu, X. I., Liu, L., Douthe, C., Li, Y., & Peng, S., et al. (2016). Rapid responses of mesophyll conductance to changes of co2 concentration, temperature and irradiance are affected by n supplements in rice. Plant Cell & Environment, 38(12), 2541–2550.

Xu, H. X. (2010). Characteristics of photosynthesis and functions of the water-water cycle in rice (oryza sativa) leaves in response to potassium deficiency. Physiologia Plantarum, 131(4), 614–621.

Zhang, W., Zhang, N. S., Zhao, J. J., Guo, Y. P., Zhao, Z. Y., & Mei, L. X. (2017). Potassium fertilization improves apple fruit (malus domestica borkh. cv. fuji) development by regulating trehalose metabolism. Journal of Horticultural Science & Biotechnology, 1–11.

Zhao, D., Oosterhuis, D. M., & Bednarz, C. W. (2001). Influence of potassium deficiency on photosynthesis, chlorophyll content, and chloroplast ultrastructure of cotton plants. Photosynthetica, 39(1), 103–109.

